# Tackling misinformation in agriculture

**DOI:** 10.1101/2019.12.27.889279

**Authors:** Jacqueline L. Stroud

## Abstract

Farmers are encouraged to embrace digital media to fill the voids caused by the privatisation of Agricultural Knowledge and Information Systems. Widespread sustainable agriculture misinformation undermines the role of science, participatory research, and evidence-based decision making. Simply providing information is insufficient, misinformation is tackled by creating a network that fosters accurate information exchange. Here I used Twitter and blended learning technologies to create a research partnership with farmers based on their beliefs that earthworms indicate good soils management. Through co-design, farmers transformed this symbol into a systematic field observation network, assessing earthworm populations to the ecological group level. Our community (#WorldWormWeek) revealed the falsehoods in misinformation such as: “Farmers around the world have been turning their fields into subterranean deserts”. This social learning network was resilient to further misinformation by the national press. Real data trends were fundamentally different to predictions made by science advancing models of global earthworm populations. Anecic earthworms (including middens) were absent in 1 in 5 fields, directly informing management practices to avoid soil biological pitfalls in no-tillage adoption. Simplistic earthworm counts to indicate soil health are rendered obsolete, a depth of information exchange can be achieved by building science-farmer partnerships using digital communications and co-designed frameworks. However, the scientific consensus, whilst generally positive about the research impact, revealed 42 % scientists rated this research as “not at all useful” or “slightly useful” to scientists. This reveals the hopeless situation where the co-production of knowledge and feedback loop linking farming-science is not broadly considered ‘science advancing’, and brought #Wormscience to an end. The next step would have been to optimize *Lumbricus terrestris* biocontrol actions targeting the soil-borne crop pathogen *Fusarium* spp. and detoxification of its mycotoxins, to reduce fungicide dependency in staple crop production; aligned with societal sustainable agriculture aspirations.

## Introduction

Sustainable agriculture relies on well-informed farmers, but there are long standing weak feedback loops between farming and science (1). Policy has delivered a global contraction of agricultural science (2), a privatization trajectory of agricultural advisory services (3), and political-economy policies that are poorly aligned to the social psychology of agro-environmental transitions (4). Globally, farmers are encouraged to embrace digital media to fill the voids caused by dismantling Agricultural Knowledge and Information Systems (5). This requires an effective discernment of information from misinformation because the latter impacts memory, reasoning and decision making, even after correction (6). Misinformation includes thought leaders selling solutions for what they describe as the ‘demineralization of our farming soils and the chemical sterilization of the soil biology’ (7), eroding trust in agricultural science and the ability of farmers to make informed decisions. Fabricated relationships between research institutes and the pesticide industry underpin conspiracy theories to question the integrity of soil scientists. A particular example is the re-interpretation of open scientific data on farmland earthworms with statements such as: “…from its earliest days, chemical farming as at Rothamsted has been detrimental to soil life and certainly is responsible for killing the soil” (8). This 2018 peer-reviewed publication has accumulated 12,000 views around the world. It was amplified into a ‘coming worm apocalypse’ using digital media platforms, and reached a global audience of millions with the message that: “Farmers around the world have been turning their fields into subterranean deserts” (9). This created tensions between scientist-farming communities, directly jeopardizing participatory soil science which is dependent on trust between these communities (10).

Participatory science/co-learning is important for developing sustainable agricultural management practices (11). These approaches link knowledge-practice-beliefs, with soils recognized as part of farmers’ identities, for example, the believe that earthworms are symbols of soil fertility and indicators of good soils management (12). However, policy makers aim to integrate nature into the economy using payment for ecosystem services constructs (13). Earthworms have been rejected from this construct for not meeting the universality required for centralized decision making by policy makers (14). Models inform policy makers of the centralized metrics to impose, provoking scientific concern about realism (15); and there are concerns that the absence of real data is no barrier for publishing payments for ecosystems services models (16). Whilst these science advances are made, farmer-driven initiatives such as Regenerative/Conservation Agriculture have created ‘anticipate no-tillage adoption’ scenarios for policy makers across Europe (17). Farmers initially use no-tillage to reduce the costs of crop establishment (17), but the process of no-tillage adoption leads to the reconstruction of their self-identity (18), and no-tillage use is reinforced by perceived environmental improvements including earthworm populations, soil carbon, soil resistance to erosion and water infiltration; often regardless of productivity outcomes (17). These farmer-driven initiatives change the role of science to a participatory role (19), which is why, for example: “Modern farming practices have killed off four out of five worms that once lived on farms” (Fig. 1) is a serious misinformation problem. Whilst soil science shows these misinformation narratives to be untrue (20), simply providing information does not solve misinformation, and is counter-productive if it challenges beliefs (6), for example, the sustainable agriculture transformation agenda. Misinformation is best tackled by the ‘pro-active formation of trusted peer networks to foster accurate information circulation for evidence-based decisions’ (6), and UK farmers indicated a potential role for scientists is to join their social learning networks for sustainable soils management (19). However, developing science-farmer partnerships is inimical to science career viability (1, 21), whilst neoliberal policies have eroded national capacities in agricultural science and information exchange mechanisms (22, 23). There is therefore no precedent on how to create a trusted peer network to foster accurate information exchange and tackle the misinformation threat to sustainable food production.

**Fig. 1.**
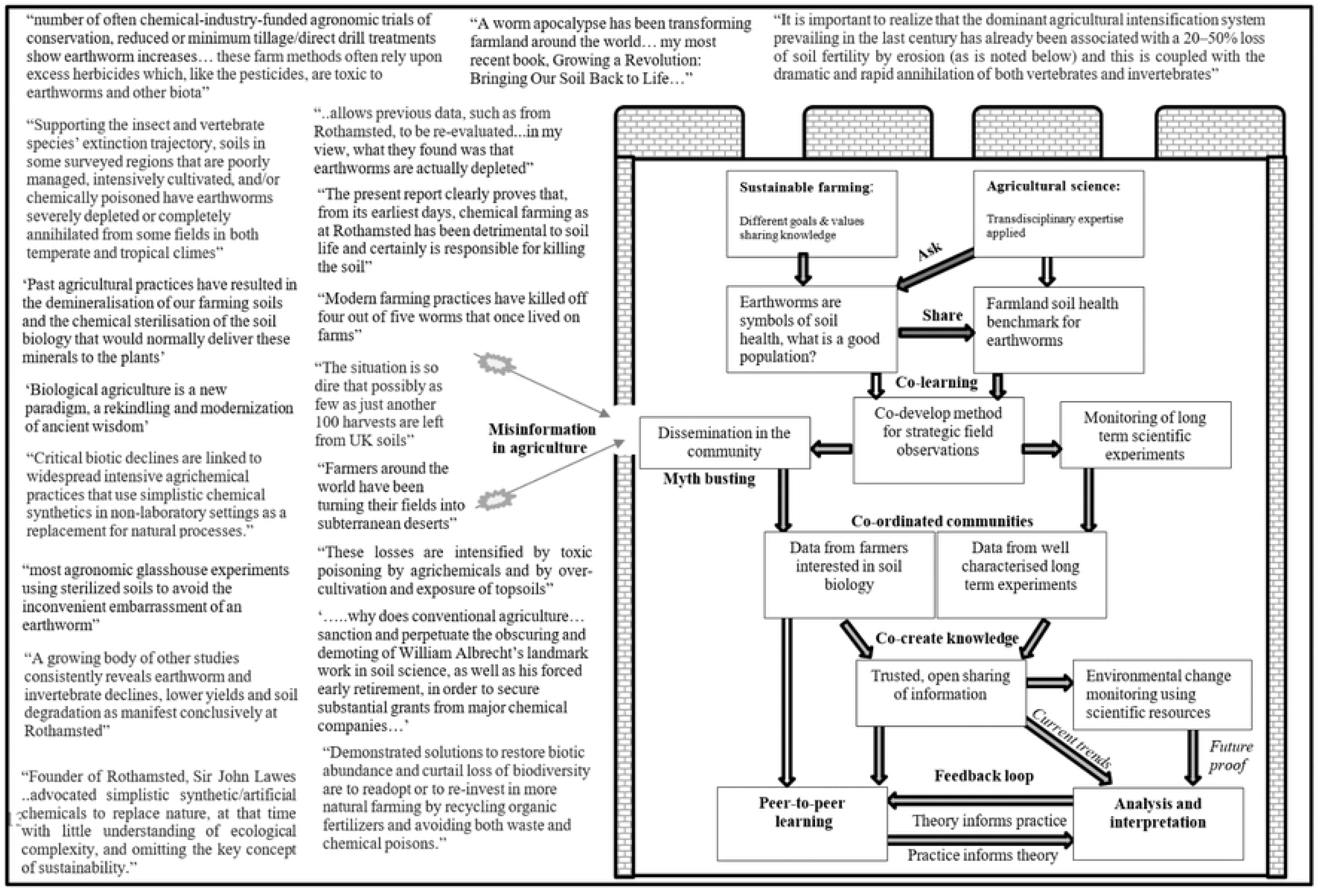
Information architecture strategy to engender resilience to misinformation (7, 8, 9).

This paper advances doing science differently through the development of a co-learning initiative within a digital information architecture strategy resilient to misinformation (Fig. 1). This is the first transdisciplinary initiative in farmland soil health that has partnered with farmers to transform a symbol into science, with the creation of a systematic field observation network, improving the ability of people to make informed decisions about soils management.

## Methods

This approach defined earthworms in a presence-absence occupancy context for farmland soil health:

- widespread earthworm presence (80+ %) within the cropped area of a field
- community structure present (epigeic, endogeic and anecic earthworm groups); linked to conventional tillage being detrimental to the two litter-feeding groups (24)

This pass/fail benchmark is a measurable earthworm configuration that enables a comparison of tillage systems and supports peer-to-peer learning. Twitter was used because of its popularity amongst farmers (25), and the network topography shifts from a centralised network that is efficient at spreading agricultural information to a decentralised diffusion system supporting learning and community cohesion (26). The structure of agricultural communities on Twitter is aligned to traditional extension services, with technical experts as ‘centralised’ hubs (here established as @wormscience). This network was managed by tagging (@) influencers to facilitate dissemination, and hashtag (#) tools connect communities to information (26).

This social learning node was integrated into the network using three steps:

a. Pilot study (#60minworms): participatory science to create partnerships with farmers and inform co-development (20);
b. Co-development phase (#30minworms): blended learning (invited face-to-face farm events and digital media technologies) to form a trusted network and test the co-designed method,
c. Application (#WorldWormWeek): to co-create knowledge and the trusted feedback loop between farming and science.

The #60minworms pilot study (29) described the principal methodology, including feedback on method compliance and essential co-developments. This was a paper-based study with booklets physically posted to participants to avoid an untested and non-peer reviewed method becoming widely available, but the role of Twitter was revealed as key to participation. Hence, this project evolved into an open digital science project, using a Twitter hub @wormscience, with # navigation to different project aspects. Twitter was also used to share research papers and observations I made from Rothamsted Research farm such as *L.terrestris* midden building, earthworm cocoon hatching, aestivating earthworms, and disservices by earthworms such as herbivory by *L.terrestris* at Rothamsted Research farm to support earthworm knowledge. I accepted all invitations to farming events during this project and each event is listed on the Twitter feed, and often includes public comments by attendees. In terms of the blended learning, I estimate I spoke to approximately 2500 people about farmland earthworms and have 3000 followers @wormscience on Twitter.

### #30minworms (September 2018 – November 2018)

I created a website for the co-development phase (www.wormscience.org), with free access to learning resources. Briefly, I made including an earthworm identification tutorial (using Powerpoint), YouTube demonstrations, the downloadable method booklet including identification guide and a link to the online form for results entry and survey feedback. I created the method booklet using free graphic design software (Canva). The sampling booklet included a 9-step method and included photos I had taken of the to the different earthworm types (epigeic, endogeic, anecic). Feedback from farmers was that the term ‘epigeic’ was too obscure and needed to be simpler, so I used: surface = epigeic, topsoil = endogeic, deep burrowing = anecic; terms to describe the different earthworm types.

Procedure: 5 soil pits per field using standard W shape field sampling

1. Dig out a 20 cm x 20 x 20 soil pit and place on soil mat (30 secs)
2. Hand-sort soil (5-mins), placing each whole worm into the pot. Record if pencil size vertical burrows are present
3. Record the total number of earthworms
4. Separate worms into adults and return juveniles to soil pit
5. Record the number of each type of adult worm
6. Return worms to the soil pit and back-fill with soil
7. Record the presence of middens
8. Repeat steps 1 - 7, until 5 soil pits assessed
9. Please input your results at www.wormscience.org for results analysis

Through uploading their results on a google form, informed consent was provided, and no identifying information was requested or recorded (i.e. the survey was anonymous). There were optional questions to provide me with feedback on the co-designed method and online training tools (Supplementary data). To ‘pass’ the test, earthworms were present in at least 4 of the 5 soil pits (widespread) and at least one adult earthworm from each ecological group (or presence of middens for anecic earthworms) to indicate earthworm diversity (all three ecological types of earthworms) are present in the field.

### Scientific consensus

There are two problems with co-designed methods, firstly the risk of bad science – peer review is too slow for this participatory process, and secondly, a risk of the sunk cost effect, is anyone going to use it and are they going to share their results? I waited until the survey met my minimum threshold of participation (50 fields, the equivalent of one week of my labour) before I emailed 30 leading scientists with publications in soils, agriculture or ecology and asked them individually to anonymously generate a scientific consensus on the co-designed method and its applications. Adopting peer-review principles, I had no collaborations with those emailed. All invitees were emailed with details of the research project website and a summary of the results (including images), and the link to the Google form which had the 20 voluntary questions and space for specific comments. Through participation, informed consent was provided. There were three compulsory questions to estimate self-selection bias linked to experience, interests and familiarity with the survey. The survey was open for 1 week and the data populated a spreadsheet (Supplementary information).

#### Dealing with misunderstanding/misinformation

The pilot study publication was reported by the national press on the 23/02/19 with ‘farmers killing soils’ headlines, with print media coverage and live BBC radio interviews. I spoke directly with reporters and journalists to try to mitigate this impact, but farmers complained to the National Farmers Union, and the NFU withdrew their support (i.e. advertising of upcoming #WorldWormWeek to their members). Hence, I decided to launch #WorldWormWeek using just @wormscience and see if there was any participation. If there was participation, I was interested to know if people would continue to report both depleted and exceptional populations (i.e. trust remained) or whether there would be a shift to just exceptional populations (politicize the survey). I emailed soil scientists around the world (China, Nigeria, Brazil, USA, Canada, Uruguay, Spain and Bangladesh) to see if there was any interest in trying the method, specifically to help create a portfolio of earthworm images, principally for machine learning applications to digitize in-field earthworm analysis for farmers, whilst creating a literal global snapshot of earthworm populations.

### #WorldWormWeek (23 – 31^st^ March 2019)

I updated the website with improved YouTube demonstrations, information about earthworms and a new booklet based on the co-development feedback. I advertised the survey using Twitter (@wormscience) and the independent dissemination of the #WorldWormWeek hashtag and ‘how to sample’ video views noted. The method booklet was principally the same, but included midden identification:

Procedure: 5 soil pits per field using standard W shape field sampling

1. At each soil pit spot, check the soil surface for the presence of middens (key shown) and tick/cross on the results sheet.
2. Dig out a 20 cm x 20 cm x 20 cm soil pit and place soil on mat (30 sec). 20 cm = 8 inches)
3. Hand-sort soil (5-minutes), placing each whole earthworm into the pot. Note if pencil size vertical burrows are present and tick/cross on the results sheet. Optional: Please take a photo of the earthworms using the image analysis sheet (scale bar & colour correction). Please upload your photos at www.wormscience.org for analysis.
4. Count the total number (adults and juveniles) of earthworms and write down
5. For UK participants: select the adult earthworms (usually only a few) and return juveniles to soil pit. Only adults have a saddle - the reproductive ring near the head. Top tip: a saddle can be more obvious if the earthworm is turned upside-down.
6. Count the numbers of each type of adult earthworm (key shown) and write down.
7. Return worms to soil pit and back fill with soil
8. Repeat steps 1 - 7, until 5 soil pits per field have been assessed
9. Please input your data at www.wormscience.org for results analysis.

The online entry portal was produced by a software company (Agrantec.com) to provide participants with an instant results dashboard with each individual results summary, and graphs of all the real-time earthworm data including numbers of earthworms and results entries submitted to facilitate peer-to-peer information exchange. The benchmark result was instantly calculated – for a field to ‘pass’ the test, earthworms were present in at least 4 of the 5 soil pits (widespread) and at least one adult earthworm from each ecological group (or presence of middens/large burrows, for anecic earthworms) to indicate earthworm diversity (all three ecological types of earthworms) are present in the field. If these conditions were not satisfied, the results were reported as a ‘fail’, and a link to a downloadable interpretation booklet was provided. Results entry included basic field management information and optional postcode for regional and soil analyses, and by using the results entry form, informed consent was provided. The photographs generated were sent to researchers specializing in machine learning technologies to try to develop digital earthworm ecological group identification tools.

### #WorldWormWeek at Rothamsted Research

The 330 ha Rothamsted Research Farm (51.82 N and 0.37 W), Harpenden, UK has a temperate climate. The soil is characterised as a flinty silty clay loam of the Batcombe series and includes the longest running agricultural field experiment in the World, with full and open experimental details available from the electronic Rothamsted Archive (www.era.rothamsted.ac.uk). I surveyed every accessible field using the #WorldWormWeek co-designed method. Modifications were made for the Fosters field experiment, a conventionally cultivated organic matter experiment with four replicate plots, so one plot was randomly sampled twice to generate five data points for each treatment, and the data averaged for ‘Fosters Field’. Individual plots (n = 9 plots, sections 0, 1 and 6, plots 2.1, 3 and 18) were sampled on the Long-Term Experiment on Broadbalk field with 5 replicate pits per plot, with the data averaged for ‘Broadbalk’.

### Open-access data sets (meteorological and soil properties)

Meteorological data (rainfall and soil temperature at 20 cm depth) was obtained for the 23 – 31^st^ March from the electronic Rothamsted Archive (www.era.rothamsted.ac.uk/Met, 2019). Regional meteorological data was obtained from the Meteorological Office MetData web application (http://basil/metdata/, 2019). Postcodes of sample locations were used to identify regional location for the meteorological context and soil properties from the UK Soil Observatory from the British Geological Society web application (http://mapapps2.bgs.ac.uk/ukso/home.html). These data were used to populate the spreadsheet, whilst maintain anonymity of the participants.

### Statistical analysis

Genstat (18^th^ addition, VSN International Ltd., UK) was used to perform t-tests to compare earthworm abundance*soil health test and earthworm abundance*tillage, with differences obtained at levels *p* ≤ 0.05 were reported as significant. The self-selection bias violates statistical assumptions for standard consolidation based data analyses (Table 1).

**Table 1.**
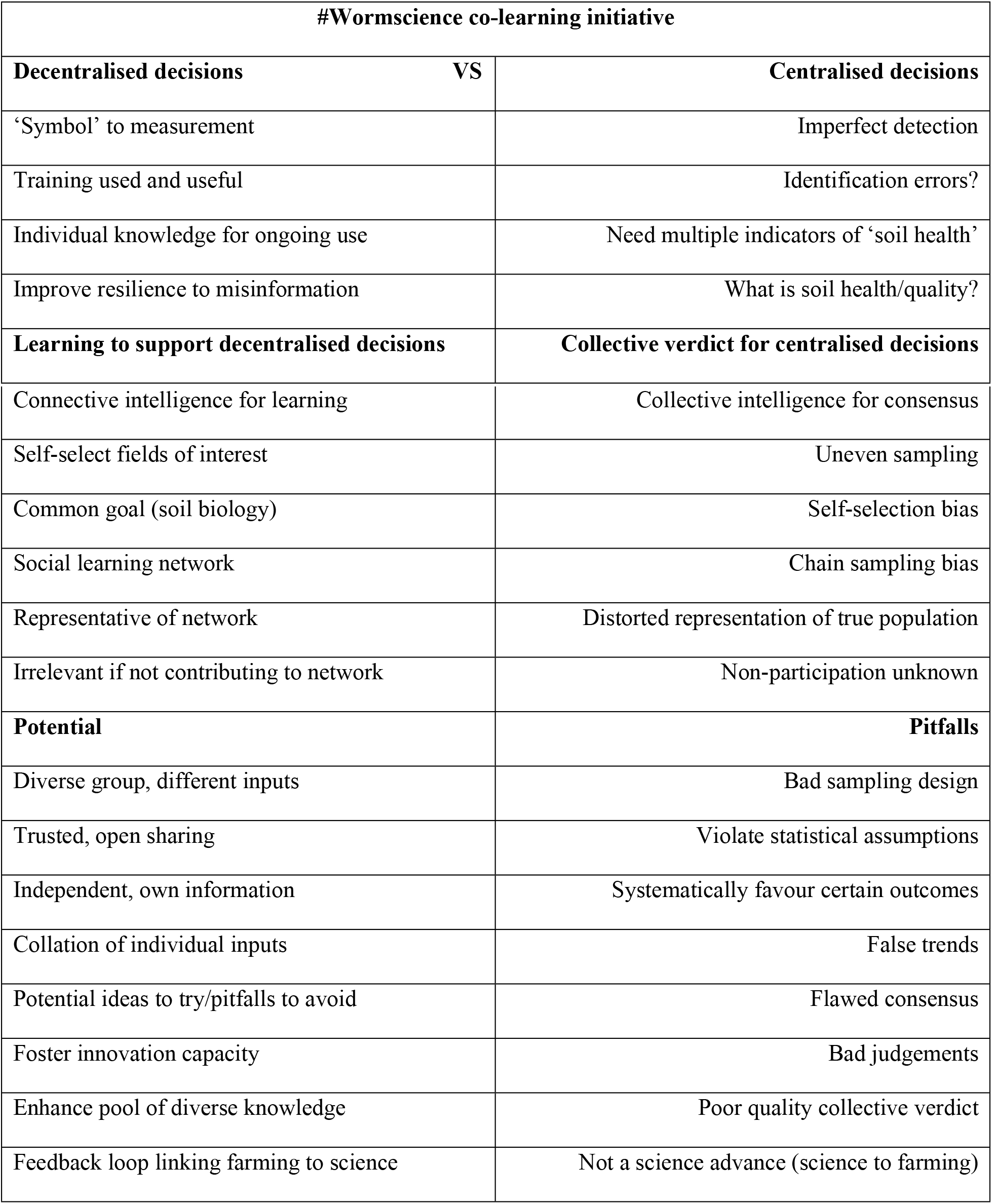
Decentralised decisions working within farmers social learning networks is a poor fit 290 for standard science approaches for centralized decisions to inform policy.

| #Wormscience co-learning initiative |  |  |
| --- | --- | --- |
| Decentralised decisions | VS | Centralised decisions |
| ‘Symbol’ to measurement |  | Imperfect detection |
| Training used and useful |  | Identification errors? |
| Individual knowledge for ongoing use |  | Need multiple indicators of ‘soil health’ |
| Improve resilience to misinformation |  | What is soil health/quality? |
| Learning to support decentralised decisions |  | Collective verdict for centralised decisions |

**Table 1.** Decentralised decisions working within farmers social learning networks is a poor fit for standard science approaches for centralized decisions to inform policy.
|  |  |
| --- | --- |
| Connective intelligence for learning | Collective intelligence for consensus |
| Self-select fields of interest | Uneven sampling |
| Common goal (soil biology) | Self-selection bias |
| Social learning network | Chain sampling bias |
| Representative of network | Distorted representation of true population |
| Irrelevant if not contributing to network | Non-participation unknown |
| **Potential** | **Pitfalls** |
| Diverse group, different inputs | Bad sampling design |
| Trusted, open sharing | Violate statistical assumptions |
| Independent, own information | Systematically favour certain outcomes |
| Collation of individual inputs | False trends |
| Potential ideas to try/pitfalls to avoid | Flawed consensus |
| Foster innovation capacity | Bad judgements |
| Enhance pool of diverse knowledge | Poor quality collective verdict |
| Feedback loop linking farming to science | Not a science advance (science to farming) |

## Results and discussion

### Scientific consensus

Changes to the pilot study included creating a range of freely available digital learning aids (www.wormscience.org). A scientific consensus approach was used, aligned with peer-review principles (30 scientists invited, 40 % engagement) to check the technical quality of the co-designed method because of the rapid development process. Experts could anonymously choose: “The survey should be discontinued due to major flaws”, but none selected this option, indicating the technical quality was sufficient. The expert consensus was that the results were likely to be caused by management practices (81 %) and that they would recommend this activity to farmers (83 %) and policy makers (58 %); 82 % thought the activity was likely to help to realize soil health in practice. However, 42 % perceived the earthworm survey as “not at all” or “slightly” useful to scientists, suggesting persistent tensions in the valuation of co-produced knowledge. Whilst the self-selection bias violates statistical assumptions for centralised decision making (Table 1); agro-environmental transformations are individual decisions informed by farmers social learning networks (18), for which this approach is well-aligned (Table 1). Whilst, this approach creates a feedback loop between farming-science by aligning with meeting farmers’ information needs, it is not aligned to science for scientists, that is, high impact papers that determine career viability (21).

### Co-development phase

Leveraging the general popularity of earthworms, ‘#30minworms’ was advertised on UK media: BBC Farming Today (listening audience, 1 million), Farmers Weekly (circulation, 53 000) and Twitter (2000 followers). Through this, farmers transformed earthworms from a ‘symbol of soil health’ into a systematic field observation network. The co-creation of knowledge attracted new co-producers (43 %) and pilot study participants (55 %), who shared the belief that earthworm monitoring is likely to improve soil health (95 %). Farmers were empowered to pro-actively educate themselves about earthworm identification using the online resources; the feedback was that this was ‘used and useful’ (91 %). They invested time (92 % completing the field survey in under a hour, aligned with method co-development improvements (20)) and physical labour to systematically measure their earthworm populations at field scales, resulting in 4647 earthworms assessed in 1628 hectares. A diverse group using different tillage practices shared their results regardless of the explicit ‘pass or fail’ soil health benchmark, fostering information exchange rather than using the survey to amplify specific beliefs. This evidences the implicit trust and/or the tolerance of an imposed soil health benchmark that underpins peer-to-peer learning (which was the principal reason for taking part (20)).

### Application phase

The publication of the pilot study (20) led to national radio and press attention, e.g. (27), but was used to spread misinformation about: ‘pesticides and farming wiping out earthworms’ just prior to the application phase. This could have ended the project by causing no further engagement or an adverse reaction. The resilience of the social learning network to this situation was tested with the launch of #WorldWormWeek on Twitter. Instead the ‘How to sample’ video was viewed 6779 times, demonstrating a strong connection to those interested in the co-creation of knowledge. The use of a hashtag #WorldWormWeek’ was effective, with its independent dissemination by national organizations including BBC Spring Watch, Woodland Trust, Natural England, Soil Association, Buglife, AHDB and DefraSoils to a total social media audience of 1 million people. The *European Journal of Soil Science* directly contributed to information circulation by launching an online special issue of open access earthworm papers. A resilient information architecture design was evidenced by the #WorldWormWeek participation increase. Based on the 10-week Autumn 2018 participation levels, a 250 % growth was needed to generate 50 entries for #WorldWormWeek. A total of 216 surveys were received, which is a growth rate of 1250 %

#WorldWormWeek resulted in a globally coordinated community of people using the same co-developed method at the same time (23 – 31^st^ March 2019) to sample farmland for earthworms (Fig. 2A, B). In the UK, both organic and conventional farmers, two groups perceived to be polarized, shared information about 9755 earthworms living in 2000 hectares of their land, enhancing the diversity of connections within the network and bringing innovators together (Fig 2 C, E). Highest global abundances were recorded in Scotland peaking at 1085 earthworms m^2^, with a UK average of 262 earthworms m^2^. This contrasts to high impact research advances, the global earthworm biodiversity model predicted abundances of 5 – 150 earthworms m^2^, (28) including Scotland having fewer earthworms than England, in agreement with a previous European model (28, 29). Differences between real data and predicted trends can be partly explained by the absence of data from England in one model (29), but both models have seemingly limited functionality for predicting biodiversity or environmental change. This suggests that science advances can be made by linking modelling to applied science, with ground-truthing improving the reliability of biodiversity predictions.

**Fig. 2.**
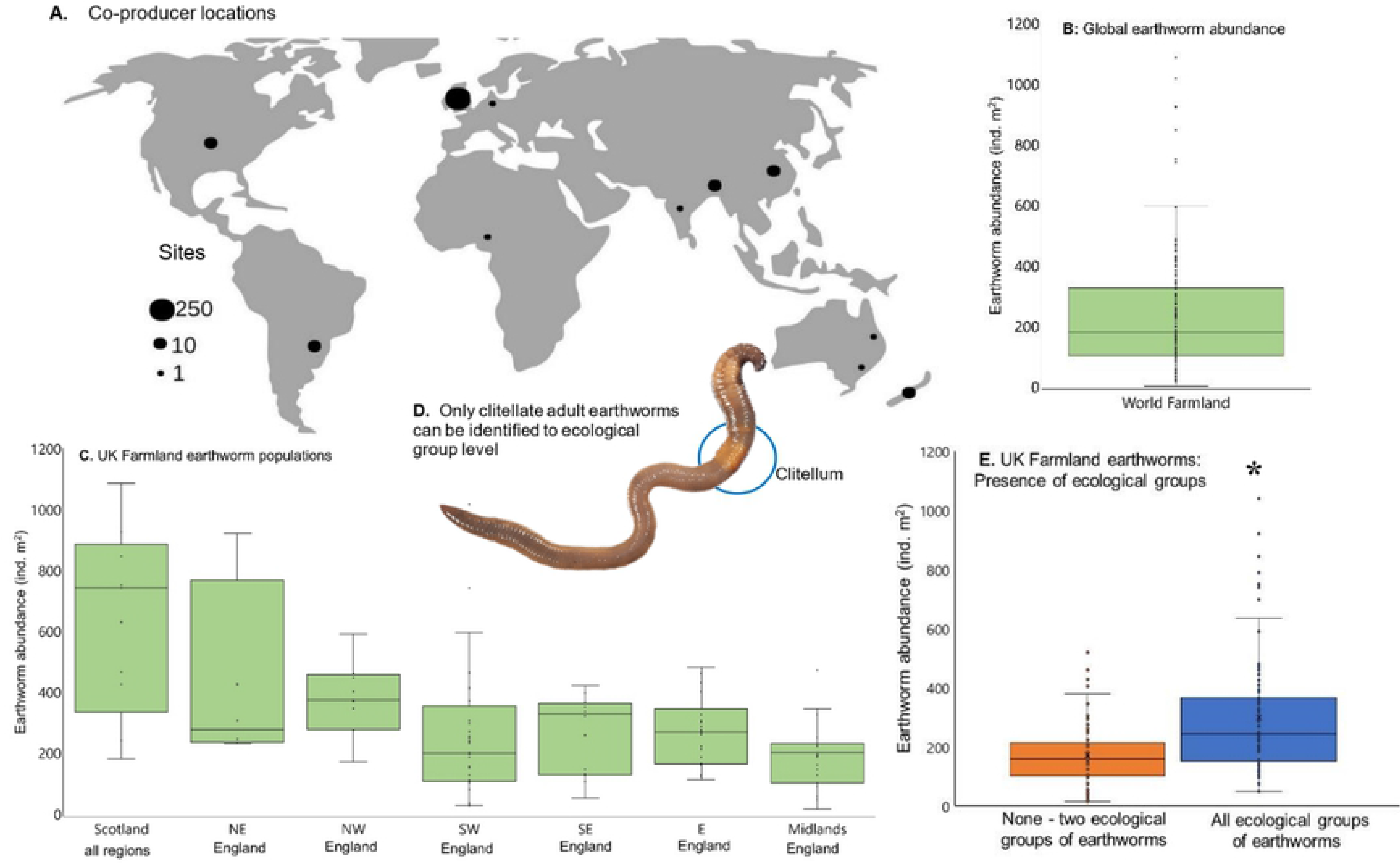
Farmland earthworm populations during #WorldWormWeek (23 – 31^st^ March 2019). A. Global participation. B. World farmland earthworm abundance; typical field abundances <150 earthworms m^2^. C. UK farmland earthworm abundances. D Ecological group analysis performed on adult earthworms. E. UK Earthworm ecological groups within fields, *significantly different.

Monitoring environmental change was trialed here through the integration of Rothamsted Farm (and its long-running experiments) into #WorldWormWeek: 86% of co-producers expressed an interest in comparing their results to the Rothamsted experiments (20). Building this connection is invaluable as open data from the long-term experiments has been used for misinformation, violating the implicit assumption that peer-reviewed scientific articles tell the truth: “…from its earliest days, chemical farming as at Rothamsted has been detrimental to soil life and certainly is responsible for killing the soil” (8). Policy makers ended research into the environmental fate and behavior of pesticides (22). In the absence of such research, this national scientific resource has been transformed into a source of global agricultural misinformation (8, 9). Such misinformation has been promoted to the extent that it has been cited in a peer-reviewed journal (30). Factually confronting such misinformation can be effective in its correction (31): earthworms are ubiquitous on Rothamsted farmland (Fig. 3). The woodland areas had fewest earthworms (0 – 60 earthworms m^2^), grass fields had 90 – 335 earthworms m^2^, and arable fields had 15 – 485 earthworms m^2^. Native habitats are often believed to have higher earthworm abundance than arable fields, and this perception is leveraged in misinformation narratives (8). Large scale Brazilian research also indicates similar trends, native forests have up to 285 earthworms m^2^, compared to farmlands with 5 – 605 earthworms m^2^ (32).

**Fig. 3.**
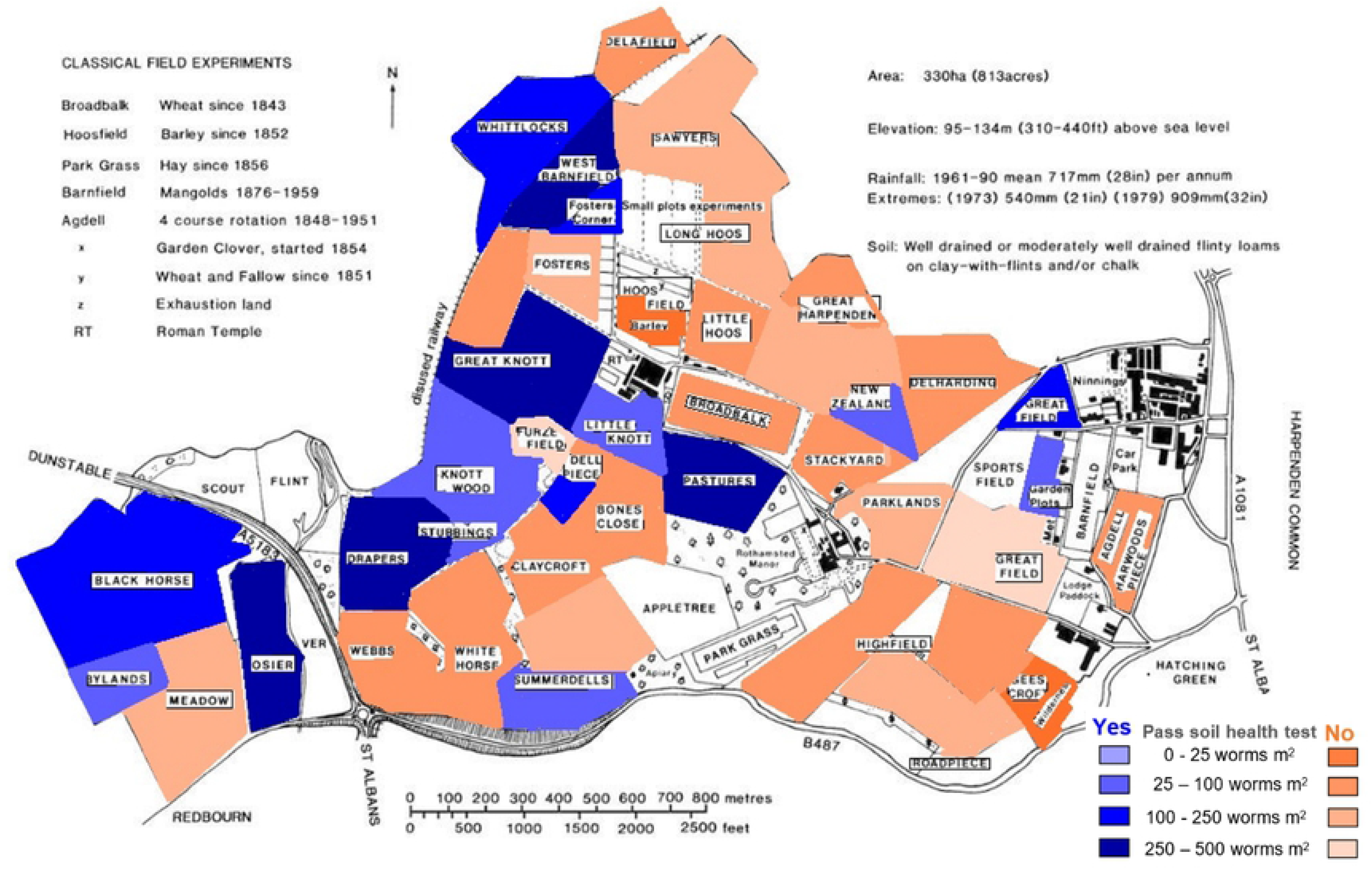
Whole farm analysis at Rothamsted Research Farm for #WorldWormWeek.

Dismantling the Agricultural Knowledge and Information System has had many impacts, for example, “research into no-tillage practices within the UK is extremely limited” (33), there are 0 public extension staff (compared to e.g. 617, 706 staff in China) (3), whilst 7 – 8 % UK farmers have adopted no-tillage, and learn from each other (34). This social learning network temporarily built a new connection between farming-science, and 61 % fields recorded the presence of adult epigeic, endogeic and anecic earthworms (Fig 4A), which was linked to the tillage system (Fig. 4B). Fields managed under no-till may have a problem from the ineffective decomposition of straw because 19 % of these fields did not detect litter-feeding epigeic earthworms (Fig. 4C). For example, a no-till co-producer reported a crop failure caused by surface straw inhibiting seedling emergence, their results showed depleted litter-feeding earthworms and informed a change in management tactics (Fig. 4C). This indicates the success of transforming a symbol of soil health, into a useful and adopted soil health measurement, in agreement with research indicating farmers use earthworms to inform management decisions (17, 20). Many tilled fields did not detect litter-feeding earthworms (epigeic or anecic earthworms) (Fig. 4D, E). An 18 % absence rate of anecic earthworms was recorded, despite the method including indicators of their activities (middens and distinctive burrows). Therefore, if farmers were to adopt no-till practices to reduce the costs of crop-establishment (17), they must avoid the common problems of poor crop emergence and a build-up of pests and diseases caused by undecomposed crop litter on the soil surface. Anecic earthworms have been successfully introduced into arable fields (35), suggesting this soil biological pitfall of no-tillage adoption is readily overcome by working with earthworm ecologists. Furthermore, environmental economists have calculated that the effective biocontrol of *Fusarium* spp. and detoxification of its mycotoxins by tillage-sensitive *Lumbricus terrestris* earthworms has an economic value of €75 ha^-1^ and would halve fungicide use (36). This research supported the development of ecological skills by farmers: identifying earthworm ecological groups, understanding the impact of tillage on earthworm community structures, and a systematic monitoring procedure to share and compare results; thus, has created a pathway to develop effective bio-control strategies in practice. This is in direct contrast to misinformation that is dependent on, and promotes ignorance: ‘no-tillage research is chemical-industry funded, it shows earthworm increases relying on excess, toxic pesticides’ (8).

**Fig. 4.**
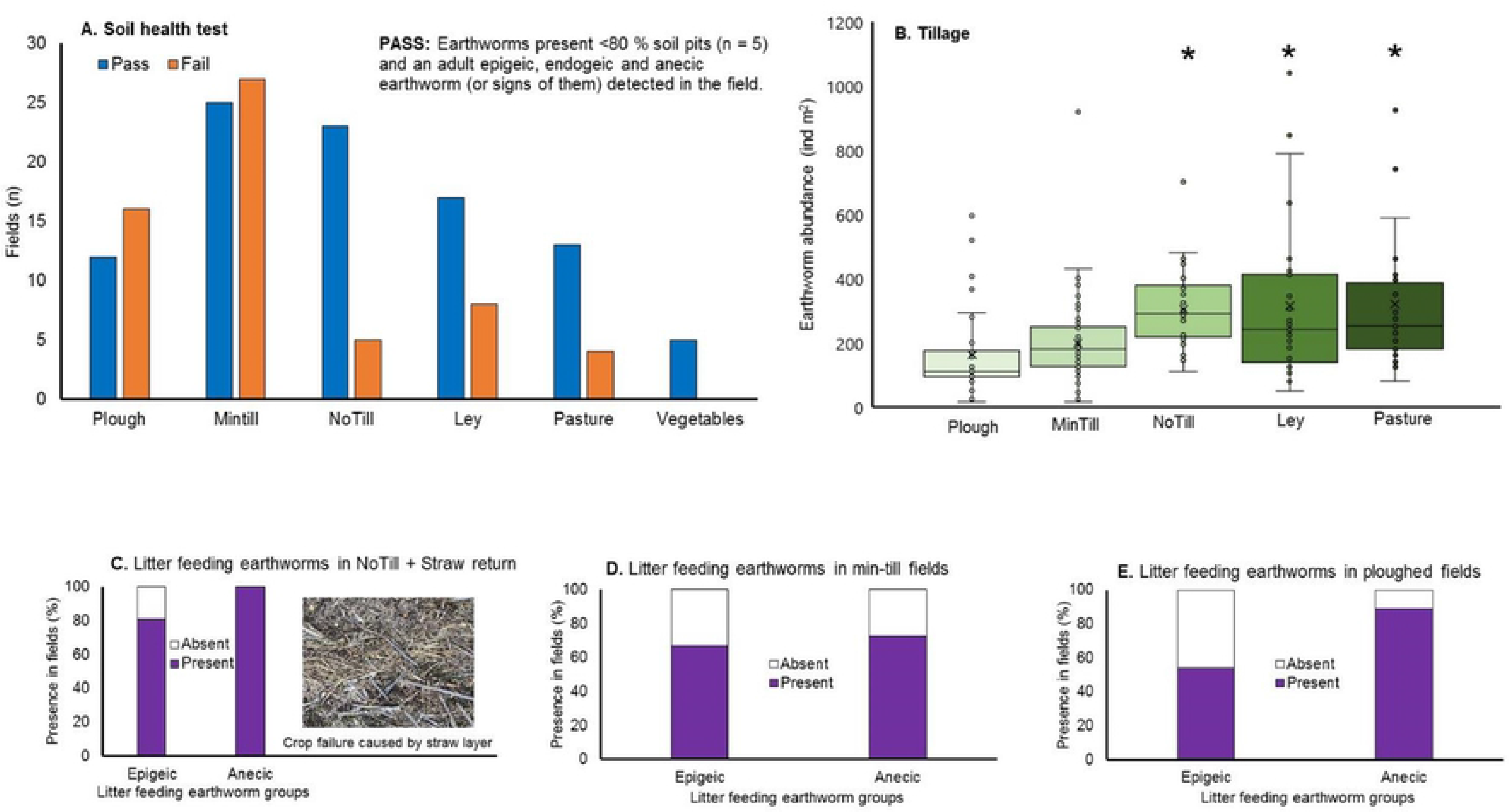
UK Farmland earthworm populations and tillage. **A**. The social learning network soil health test results. **B**. Tillage influences on earthworm abundance, * indicates significantly different. C-D: litter feeding earthworms in: **C** No-tillage with straw (photograph from a co-producer showing the straw problem); **D** MinTill; **E** Ploughed fields.

Policy promotes the perception that earthworm counts can be used to indicate soil health (37), but this simplistic metric is rendered obsolete compared to what can be achieved by building science-farmer partnerships using digital communications and co-designed frameworks. Farmers have proven capabilities in developing skills in earthworm identification, including *L.terrestris* middens. Other studies have shown that there is a positive correlation between midden abundance and arable soil health scores, linked to beneficial earthworm impacts on soil microbiology and soil physical structure (38), further highlighting the redundancy of earthworm counts. Middens can cover around 25 % of the soil surface and influence the spatial distribution of soil processes including organic matter decomposition, N-mineralisation and water movement (39). Soil organic carbon and nitrogen is stored in better protected soil fractions of clay soils in middened areas compared to non-middened areas within a field (40). These earthworm burrows have elevated levels of NO_3_^-^ and NH_4_^+^, and enriched populations of nitrifying and denitrifying bacteria than the bulk soil (41). Middened areas are associated with 43 % higher NOx emissions compared to non-middened areas of soils (42). Middens are often rare in conventionally tilled fields (43). Together this suggests the potential to develop middens as a keystone indicator of soil processes in farmers’ fields. However, participatory approaches are rarely considered to be a legitimate route for research funding in high income countries (12).

Neoliberal policies have eroded national capacities in agricultural science and information exchange mechanisms (22, 23). Yet, neoliberal policy makers directly advocate for science and technology to create a knowledge economy (44), suggesting there is an overlooked disconnection between science-policy that is inhibiting realizing a knowledge economy in practice. This research was initiated by the use of open data (that is supposed to improve the trust in science), for a misinformation narrative that went viral and tested the trust of co-producers (10), suggesting a widespread naiveite in potential open data uses. There is prevalent evidence that farming-science relationships are generally impaired; Australia, the first country experimenting with privitisation has scientists calling for public re-investment to tackle disillusionment and sustainability issues (23), whilst unprecedented hostility by the farming community towards the soil scientists has been reported (45). In the UK, there is a loss of trust in the Department of Food, Environment and Rural Affairs caused by its actions to dismantle knowledge networks (46), and hostile interactions with scientists and the farming community are shared using digital media platforms, for example: “I don’t know what the value of science is, farmers, at least in my experience in North America, have written off the science, the scientists, the universities…” (47). America is starting this privatization journey, with researchers concerned about the impact of neoliberal policies on soil sustainability (48). Modelling approaches are useful but create an illusion of simplicity aligned with simple rational-actor models of human behavior that underpin political-economy policies (13). The neglect of the complex connections between people and soils is problematic as this is key to enhancing the pro-environmental behaviours that are needed to bring about a sustainable food production system (49).

## Additional information

The author declares no competing financial interest.

## Supporting information

**Supplementary information.** All raw data is available as a spreadsheet.

